# Cerebral organoids at the air-liquid interface generate diverse nerve tracts with functional output

**DOI:** 10.1101/353151

**Authors:** Stefano L. Giandomenico, Susanna B. Mierau, George M. Gibbons, Lea M.D. Wenger, Laura Masullo, Timothy Sit, Magdalena Sutcliffe, Jerome Boulanger, Marco Tripodi, Emmanuel Derivery, Ole Paulsen, András Lakatos, Madeline A. Lancaster

## Abstract

Three-dimensional neural organoids are emerging tools with the potential for improving our understanding of human brain development and neurological disorders. Recent advances in this field have demonstrated their capacity to model neurogenesis^1,2^, neuronal migration and positioning^3,4^, and even response to sensory input^5^. However, it remains to be seen whether these tissues can model axon guidance dynamics and the formation of complex connectivity with functional neuronal output. Here, we have established a longterm air-liquid interface culture paradigm that leads to improved neuronal survival and allows for imaging of axon guidance. Over time, these cultures spontaneously form thick axon tracts capable of projecting over long distances. Axon bundles display various morphological behaviors including intracortical projection within and across the organoid, growth cone turning, decussation, and projection away from the organoid. Single-cell RNA-sequencing reveals the full repertoire of cortical neuronal identities, and retrograde labelling demonstrates these tract morphologies match the appropriate molecular identities, namely callosal and corticofugal neuron types. We show that these neurons are functionally mature, generate active networks within the organoid, and that extracortical projecting tracts innervate and activate mouse spinal cord-muscle explants. Muscle contractions can be evoked by stimulation of the organoid, while axotomy of the innervating tracts abolishes the muscle contraction response, demonstrating dependence on connection with the organoid. Overall, these results reveal a remarkable selforganization of corticofugal and callosal tracts with a functional output, providing new opportunities to examine relevant aspects of human CNS development and response to injury.

---

Cerebral organoids^1^ model early brain development with remarkable fidelity, but later neuronal maturation is limited by insufficient oxygen and nutrient availability due to the lack of a blood supply. Recent advances have been made to improve survival either by providing growth factors such as BDNF^5^ or by transplanting neural organoids into a rodent host to allow for vascularization^6^. In an effort to improve oxygen supply but retain the accessibility and scalability afforded *in vitro*, we tested whether the classic approach of organotypic slice culture^7^ at the air-liquid interface could be applied to organoids to improve long-term survival. After optimizing sectioning and culture conditions (see Methods, Figure 1a), we found that air-liquid interface cerebral organoids (hereafter referred to as ALI-COs) displayed overall improved morphology compared with whole organoids (Figure 1b) as well as increased numbers of cortical neuron populations, suggesting improved survival (Figure 1c).

**Figure 1.**
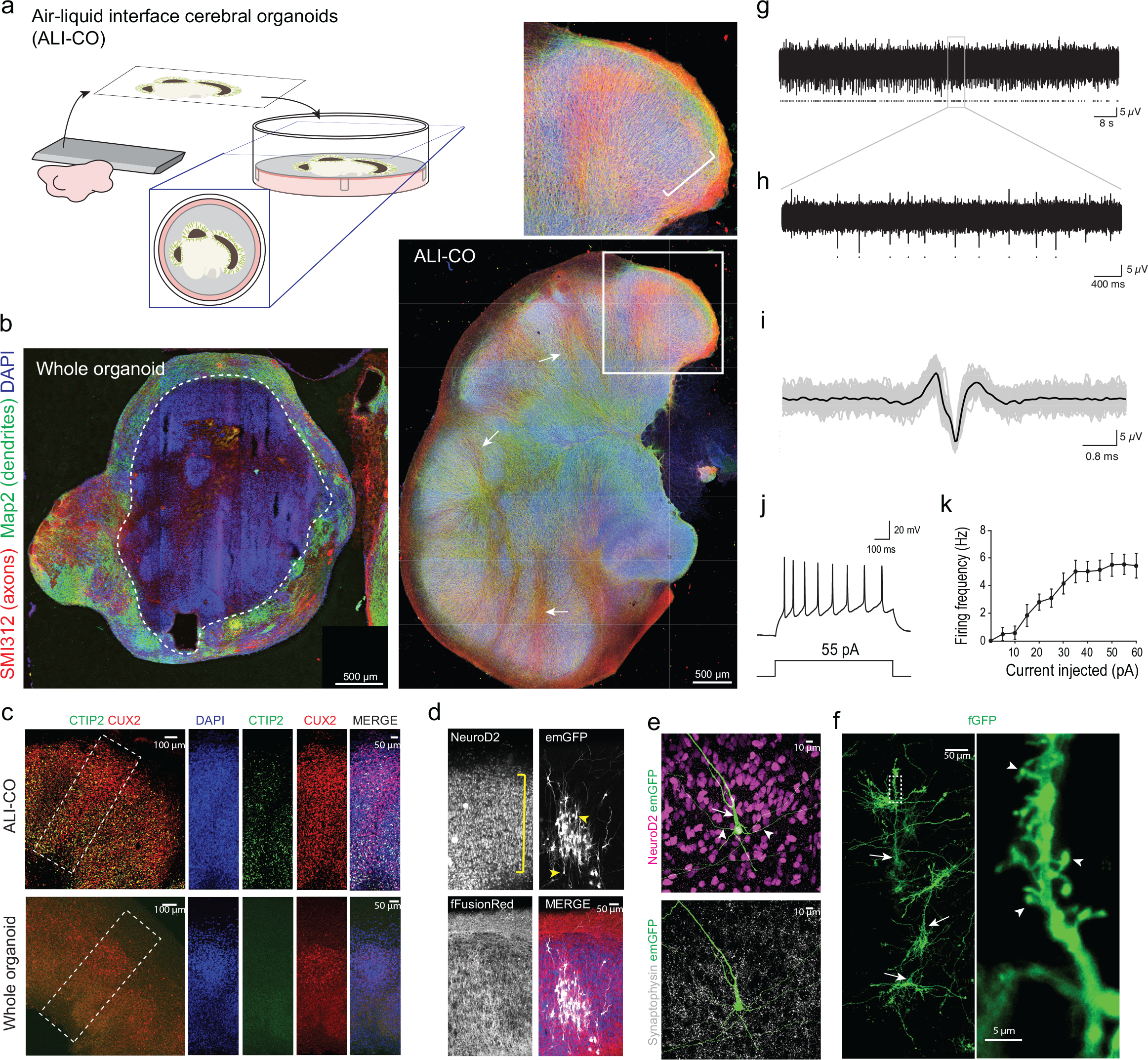
Culture at the air-liquid interface leads to improved neuronal survival and maturation. **a.** Schematic of the culture paradigm as detailed in methods. **b.** Immunohistochemistry with the marker SMI312 (red) to stain axons, and MAP2 (green) for dendrites, on representative sections of whole versus ALI cerebral organoid cultures. Both were generated using the same protocol except that the ALI-CO was sectioned at 61 days and cultured at the ALI for an additional 21 days (82 days total). Whole organoid age was 90 days. Dashed line indicates the border of healthy neurons along the organoid surface. Inset shows a higher magnification of a lobule containing radially aligned neurons of the cortical plate (bracket) and arrows indicate SMI312 positive inward projecting tracts. **c.** Staining for deep layer (Ctip2+) and upper layer (Cux2+) neurons reveals increased numbers of both populations in ALI-COs with a particularly strong effect on deep layer neurons where most of the staining for Ctip2 is unspecific in whole organoids at this stage (whole organoid age: 116 days, ALI-CO age: 84 days + 36 days ALI). **d.** Sparse labeling in a 43+37 day ALI-CO by Sendai-virus encoding emGFP (white) reveals radially aligned neurons (NeuroD2, blue) with complex dendritic architectures and pyramidal morphologies (arrowheads) within the aligned cortical plate (bracket). All cells are labeled with fFusionRed (red) to visualize the overall morphology. **e.** Higher magnification of a single emGFP (green) labeled neuron (NeuroD2+, magenta) displaying typical pyramidal morphology with primary dendrite (arrow) and basal dendrites (arrowheads). Synaptophysin (white) reveals extensive synaptic staining. **f.** Electroporation of membrane targeted farnesylated GFP (fGFP, green) reveals complex dendritic architecture of radially aligned pyramidal neurons (arrows) with evident dendritic spines (arrowheads, inset). **g.** Two minutes of spontaneous activity recorded from a single electrode of a multi-electrode array (MEA) in a day 63+54 ALI-CO; detected action potentials marked with dots. **h.** Five-seconds from the same recording (expanded from grey box). **i.** Overlay of all detected spikes from this electrode (grey, marked by dots in g) with mean waveform in black. **j.** Whole-cell patch clamp recordings of action potentials evoked by a 55pA current injection. **k.** Frequency-current (F-I) curve showing average action potential firing rate with increasing amplitude of current injection (mean+s.e.m., n=13 cells from 7 ALI-COs).

Sparse labelling with GFP (Figure 1d) revealed neurons that displayed mature morphologies (Figure 1e) with complex dendrites and dendritic spines (Figure 1f). Staining for pre- and post-synaptic markers demonstrated synapses decorating these dendrites (Supplemental Fig. 1a). We also observed various interneuron types and the presence of GABAergic synapses (Supplemental Fig. 1b-d). This suggests the various cortical neuron and synapse types are present to allow for functional circuit formation. Indeed, ALI-COs stained positive for the marker of neuronal activity c-Fos (Supplemental Fig. 1e). Multi-electrode array recordings (Supplemental Figure 1f) revealed the presence of spontaneous neural activity (Figure 1 g-i, Supplemental Fig. 1g), which was blocked by tetrodotoxin (TTX, Supplemental Fig. 1h-k). We also performed whole-cell patch clamp recordings of individual neurons demonstrating the ability to fire trains of action potentials with positive current injection (Figure 1j, k). These findings suggest the generation of mature neuronal morphology and activity in ALI-COs.

Because the slice culture is easily tractable for live imaging, we examined axon guidance in GFP-labelled neurons over long-time periods (Figure 2a, Supplemental Fig. 2a). Initial axon outgrowth was highly dynamic (Figure 2b, Supplemental Fig. 2b, Supplementary Videos 1, 2) while later axons projected in a directional fashion (Figure 2c, Supplementary Video 3) and with greater velocity within bundles of axons (Figure 2d, Supplemental Fig. 2c, Supplementary Videos 4, 5). These behaviours are reminiscent of the axon outgrowth dynamics of pioneer axons and follower axons of established tracts^8^. Indeed, ALI-COs displayed robust bundles of axons (Supplemental Fig. 2d) that became reinforced over time (Figure 2e) rather than randomly filling with axons in all directions as is more typically seen *in vitro*^9^. Overall, the resultant axon bundles showed specific orientations and a high degree of coherency that is typical of tract morphology (Figure 2f, Supplemental Figure 2e).

**Figure 2.**
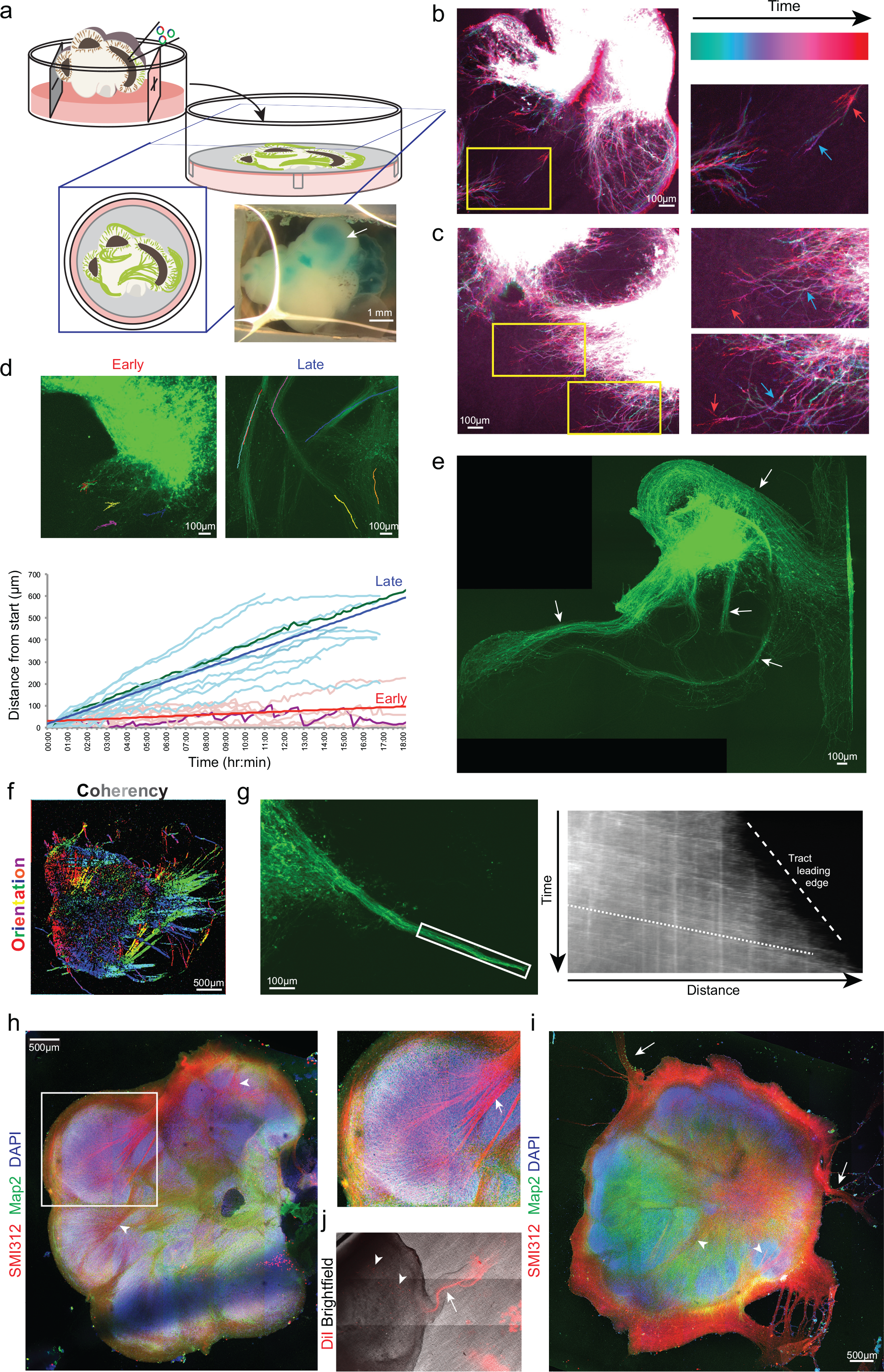
Neurons of ALI-COs exhibit dynamic axon guidance behaviors and establishment of tracts. **a.** Schematic of electroporation and preparation of ALI-COs for live imaging. Inset image shows an organoid after plasmid injection along with a blue dye (FastGreen) to visualize the injected ventricle (arrow). **b.** Temporal projection image pseudocolored by time (Supplementary Video 2) of an electroporated ALI-CO 5 days after placement at the ALI showing dynamic axon outgrowth with growth cone retraction (blue arrow to red arrow in the higher magnification inset). **c.** Temporal projection image (Supplementary Video 3) after 9 days at the ALI showing more directed axon outgrowth with progressive extension (blue arrow to red arrow in higher magnification insets). **d.** Tracing of individual growth cones over time from ALI-COs at the ALI for 2-5 days (early) and 14-24 days (late) reveals disparate behaviors and velocities (see Supplementary Videos 1, 4). Growth cones in later, established tracts exhibit a higher velocity (bold blue and red lines are average linear regression for each set of data), while earlier growth cones exhibit dynamic retractions (visible in a highlighted trace in purple compared with highlighted green trace of a later timepoint). An example image of each is shown above with superimposed traces. Tracing was done on 9 growth cones (early) and 12 growth cones (late) from four samples. **e.** Axon tracts after 18 days ALI culture demonstrating numerous dense bundles (arrows) with nonrandom projection pattern. Note the same overall pattern but with reinforced bundles compared with Supplemental Figure 2d, which was taken 4 days earlier. **f.** Directional image analysis for the orientation (color coding) and coherency (brightness) of axon tracts (original image Supplemental Fig. 2e). **g.** Still frame (left) and kymograph (right) of an extending tract (boxed region) (Supplementary Video 8). Note the higher velocity (shallow slope, dotted line) of incoming growth cones while the leading edge of the tract as a whole progresses more slowly (steep slope, dashed line). **h.** Staining for all axons (SMI312) and dendrites (Map2) on a whole ALI-CO after 36 days at the ALI reveals thick bundles (arrowheads) that can be seen projecting within the organoid and merging to form large tracts (inset, arrow). **i.** Axon (SMI312) and dendrite (Map2) staining of another ALI-CO after 41 days at the ALI reveals the presence of tracts projecting away (arrows) from the main mass in addition to intra-organoid projections (arrowheads). **j.** Retrograde labeling with DiI of an escaping tract reveals the morphology of the tract (arrow) and the location of projecting cells (arrowheads) within the organoid (Supplemental Figure 2h).

Interestingly, these axon tracts could be observed to project within local regions, across the organoid over long distances, and away from the organoid altogether (Figure 2e, Supplemental Fig. 3f). Some tracts crossed in a manner similar to decussation (Supplemental Fig. 2f, Supplementary Video 6), with incoming growth cones retaining their directionality upon arrival at the intersection, while other growth cones and tracts could be seen exhibiting turning behaviors (Supplemental Fig. 2c, g, Supplementary Video 7). Finally, axon bundles could be seen extending as a whole, with incoming growth cones projecting at a fast pace, while the tract leading edge progressed much more slowly (Figure 2g, Supplementary Video 8).

Live imaging results point to the establishment of diverse axon tract morphologies; however, these visible tracts are only a subset of axons labelled by electroporation. In order to visualize the full diversity of axon tracts, we stained mature ALI-COs with SMI312, a broad marker of axons^10^, which could be seen originating within discrete lobules and merging to form dense axon bundles (Figure 2h) reminiscent of intracortical tracts of the CNS. Furthermore, we observed outgrowth of axon tracts projecting away from the organoid (Figure 2i), and retrograde tracing with the lipophilic dye DiI (Figure 2j) better revealed the morphology of these escaping tracts as well as the location of the responsible projection neurons (Supplemental Fig. 2h).

The presence of a variety of axon tract morphologies suggests that ALI-COs might display various neuron identities that could exhibit different projection behaviors. In order to test this possibility, we performed single-cell RNAseq on ALI-COs to examine the full repertoire of cell types. We analysed six slices from three ALI-CO preparations with an average of 4,427 cells per sample, which were processed through the 10X single-cell genomics platform (see Methods). Unbiased clustering of cell populations was achieved by principle component analysis (PCA) using highly variable genes as input and was visualized following dimensionality reduction using tSNE embedding (Figure 3a). This resulted in the separation of six well-defined major clusters (C1-C6, Figure 3b).

**Figure 3.**
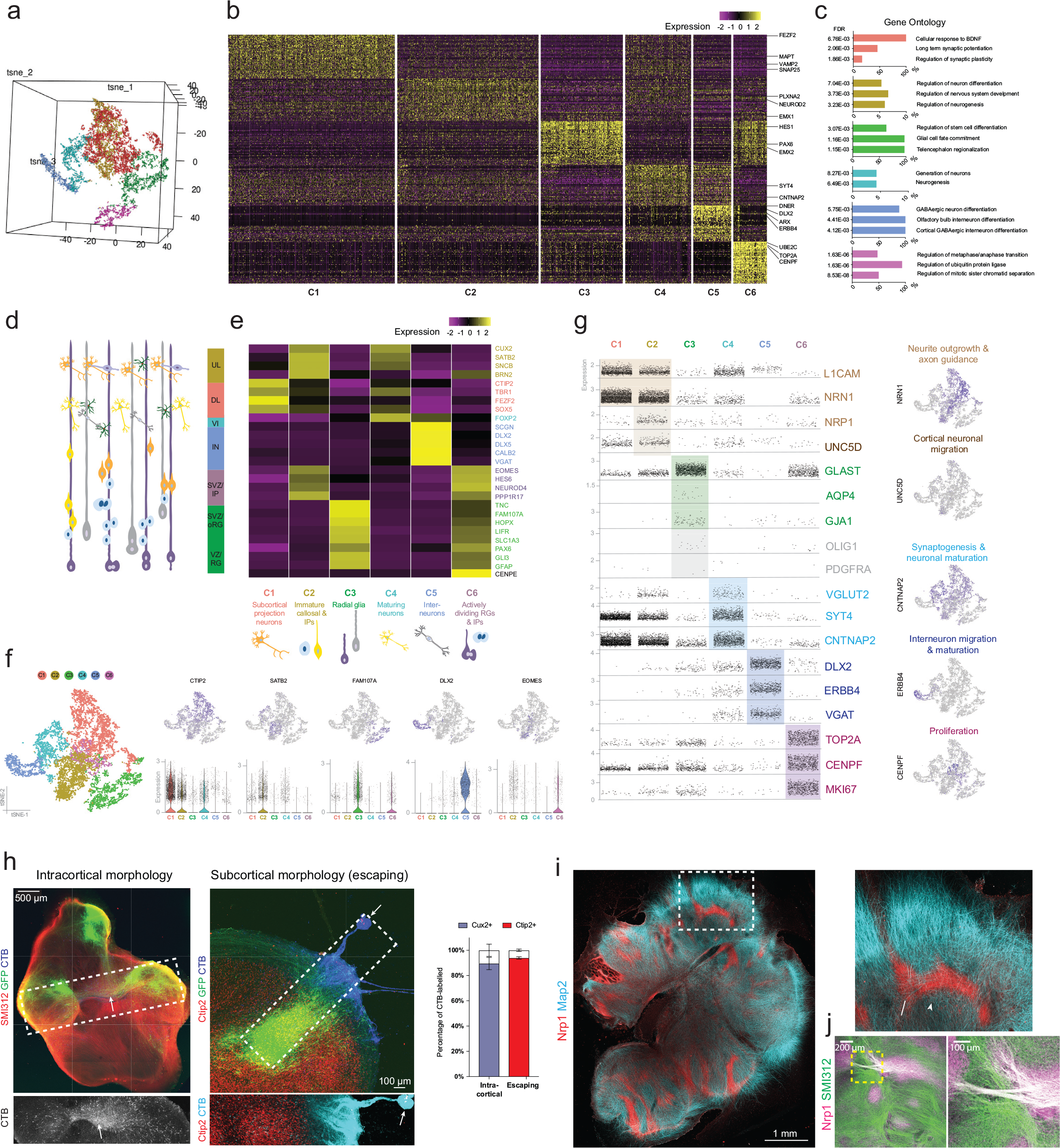
ALI-COs contain diverse neuron identities that exhibit specific projection patterns. **a.** Unbiased tSNE separation of clusters based on single-cell RNA-sequencing data visualized on a 3D space, showing six main populations. **b.** Heatmap demonstrating scaled levels of the top 50 differentially expressed genes in the main clusters (C1-6) with example genes (shown at right). **c.** Histograms of the three top gene ontology (GO) terms with the highest fold-enrichment values (FDR above 0.1%, Fisher’s test) of the top 50 genes for each population. Colour coding of bars represent cluster identities. **d.** Schematic representation of cells residing in the different layers of the fetal human cortex: ventricular zone (VZ) containing radial glia (RG), subventricular zone (SVZ) containing outer/basal radial glia (oRG) and intermediate progenitors (IP), deep (DL) and upper cortical plate layers (UL) as well as interneurons (IN) and layer 6 neurons (VI). **e.** Heatmap showing the scaled mean expression levels of layer- and cell-type specific genes within the six unbiased clusters, identifying the major cell populations. The colour coding of layers, marker genes (right side) and cell types (bottom) corresponds with cluster identities (C1-C6, bottom). **f.** 2D tSNE plots of clusters (left side) and the distribution of cells expressing layer or cell-type marker genes across the main populations (upper row). Scatter plots below show the distribution of corresponding normalized gene expression values per cell within each cluster, and the violin plot is displayed where the proportion of cells expressing a given gene is the highest. **g.** Scatter plots demonstrating the normalized gene expression levels of genes per cell across the six main clusters with a focus on relevant neuronal, progenitor and glial functions. Feature plots (right hand column) demonstrate example genes expressed by populations confined to a particular cluster or by multiple cell populations. Colour coding represent functional gene associations. **h.** Retrograde labeling by CTB microinjection (arrow indicated injection site) reveals disparate identities contributing to distinct tract morphologies. Tracts with internal projection morphology (left) traced back to primarily Cux2+ callosal projection identity cells (quantified at right) while escaping tracts traced back to primarily Ctip2+ subcortical projection identity cells. Three ALI-COs for each condition (intracortical and escaping) were labeled and all CTB-labeled cells in three z-planes of whole ALI-CO slices were counted for Cux2 or Ctip2 staining. Error bars are SEM. **i.** Staining for the marker of corpus callosum, Nrp1, reveals several internal tracts that are positive. Map2 stains dendrites. **j.** Costaining of Nrp1 and SMI312 reveals a specific subset of tracts that are positive for Nrp1 suggesting callosal identity.

GO term analysis (FDR > 0.1% with highest fold-enrichment) of the top 50 differentially expressed genes (Figure 3c) raised a possibility that developmental cell states define the main cell populations, reflecting stages of *in vivo* cortical development (Figure 3d). In support of this, gene expression correlation (Supplemental Fig. 3a) and pseudotime analyses (Supplemental Fig. 3b, c) revealed a similar co-expression pattern of progenitor zone and neuronal layer marker genes in the organoids to that shown for the age-matched fetal brain. Thus, we next compared the average expression of developing cortical cell-type-, cell state-, and region-specific marker genes (see Methods). Cluster identities better corresponded with cell-types and maturity (Figure 3e) than brain region specific markers (Supplemental Fig. 3d,e). This also supports previous findings that the enCOR method used as the starting material for ALI-COs preferentially generates a forebrain identity^4^.

Cell-type and maturity markers revealed a distinct population of deep layer subcortical projection neurons (C1: e.g. CTIP2, FEZF2), upper layer intracortical (callosal) projection neurons and intermediate progenitors (C2: e.g. SATB2, EOMES), ventricular and subventricular zone radial glial cells (C3: GFAP, FAM107A), more mature upper and deep layer neurons (C4: e.g. FOXP2, CUX2), interneurons (C5: e.g. DLX2) and actively dividing cells with intermediate progenitor and radial glia markers (C6: e.g. CENPE, EOMES, GLAST) (Figure 3d-f). Staining for upper layer and deep layer identities further supported the presence of a variety of projection types concomitant with diverse tract morphologies (Supplemental Fig. 3f). Interestingly, the overlapping cluster of upper layer neurons and intermediate progenitors suggests many of the upper layer neurons may still be quite immature, which would match previous findings that upper layer neurons are actively being generated at this time point (69-75 days) in organoid development^11^.

We then examined if the cluster forming cell types were associated with a gene expression profile that appropriately corresponds with their expected functional characteristics, such as axon-projection and neuronal circuit formation. Many of these were also present within the differentially expressed genes (Figure 3g). Genes associated with axon outgrowth/tract formation (L1CAM, NRN1) were expressed more abundantly and at increased levels in the C1-2 clusters formed by extra- and intracerebral projection neurons and to lesser extent in the C4 population. The C4 cluster showed higher levels of more mature markers of excitatory and inhibitory synapse formation (VGLUT2, CNTNAP2, SYT4). The high relative expression of DLX2, DLX5, ERBB4 and VGAT in C5 further indicated a very distinct interneuron population and, together with histological data (Extended Figure 1b-d), raised a possibility for the presence of early cortical circuits. Some cells with functionally relevant markers of astroglial maturity (AQP4, GJA1) and the oligodendrocyte-lineage (OLIG1, PDGFRA) were present in C3, but to a very low degree at this relatively early stage (C3). The C6 progenitor population displayed high levels of transcripts indicative of cell proliferation (CENPF, TOP2A), corresponding with the amplifying cortex-forming populations. Overall there was a well-defined association between cell-types and their expected function suggested by the molecular profiles in each cluster (Figure 3f, g).

To test whether, as seen *in vivo*, these upper and deep layer neuron populations in ALI-COs project as tracts with distinct morphology, retrograde labelling with cholera toxin subunit B (CTB) was performed on escaping and internal axonal bundles. Whilst 93.9% (SD=0.9, n=3) of neurons projecting into escaping tracts stained positive for the subcortical projection marker CTIP2, 89.5% (SD= 4.8, n=3) of neurons projecting into internal bundles stained positive for the callosal projection neuron marker CUX2 (Figure h). These data suggest the morphology of these tracts matches the correct molecular identity. Furthermore, staining for the specific marker of the developing corpus callosum, NRP1^12^, revealed a subset of internal projecting ALI-CO tracts (Figure 3i) that often appeared thicker and more fibrous than NRP1-negative tracts (Figure 3j).

We sought to test the functionality of these tracts, focusing first on the internal projecting tracts. We employed 3D multi-electrode arrays (MEAs) to perform extracellular recordings (Figure 4a) and infer the functional connectivity within ALI-COs from analysis of correlated spontaneous activity^13^. ALI-COs showed network bursts (Figure 4b, Supplemental Fig. 4a) in which neurons near multiple electrodes across the array of electrodes showed simultaneous bursts of action potentials, as seen in mature neuronal networks^14^. Comparison of the correlated activity revealed densely connected local networks (Figure 4c, Supplemental Fig. 4b) in which the highly correlated neuronal activity could be found within both short- and long-range connections between nodes (Figure 4d). Interestingly, while the connections appeared stronger over shorter distances, the highest correlated activity occurred at a greater distance than the distance between two electrodes (which are 200 μm apart) suggesting these are not simply nearest neighbour connections but that there is a degree of spatial specificity in the connections made. Overall, these findings point to specific spatial patterns of connectivity within ALI-CO cultures.

**Figure 4.**
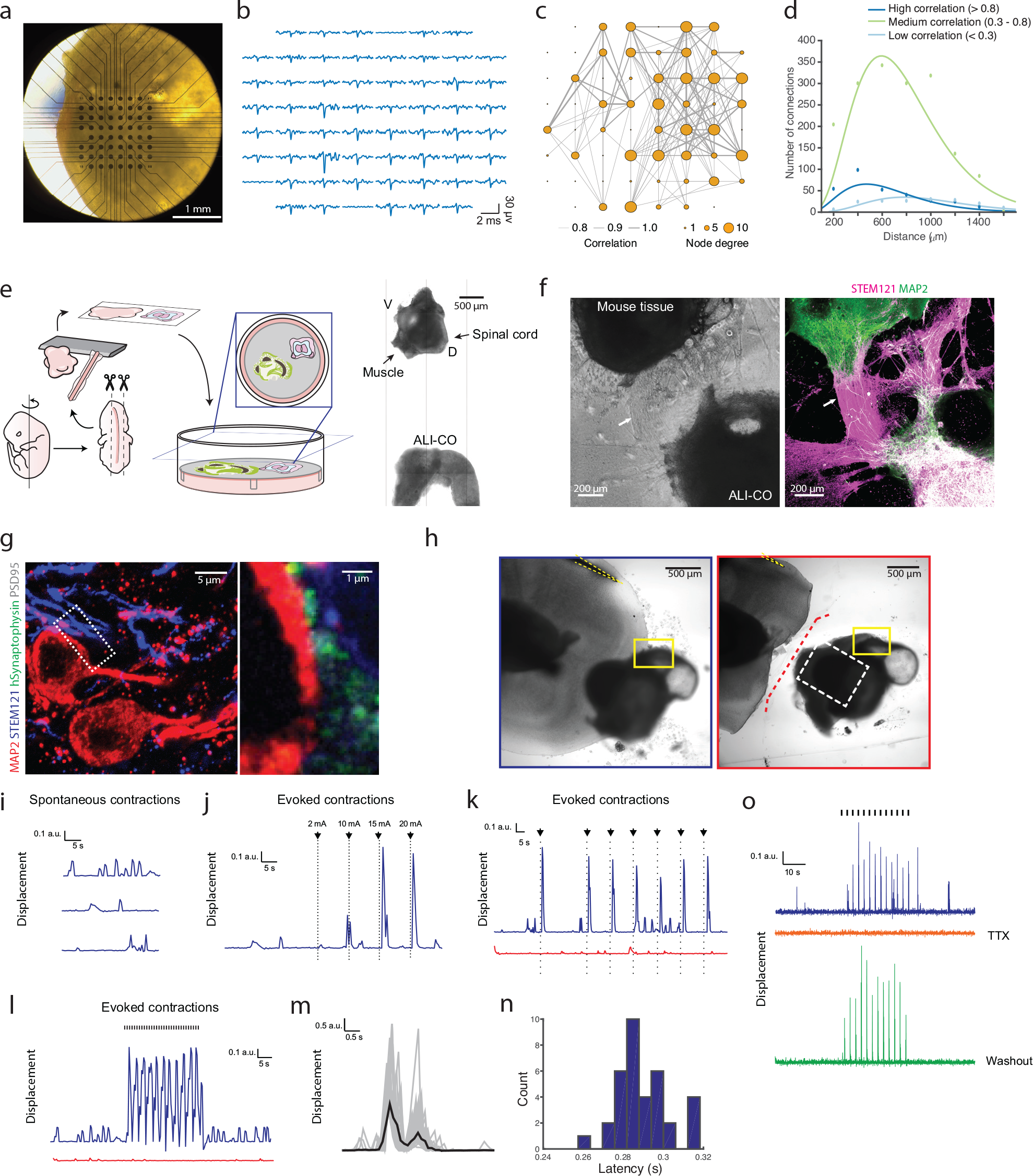
Functional connectivity in intracortical and extracortical projecting axon tracts. **a.** Photo of ALI-CO at 39 days of ALI culture upon transfer to the 3D microarray. **b.** 3.4-millisecond-long traces of spontaneous activity in the 59 electrodes at the time of a network burst. **c.** Network plot showing functional connectivity between specific sites within the ALI-CO. Line thickness represents strength of correlation in the spontaneous activity recorded between connected nodes (MEA electrodes) in the network, while size of the node indicates the number of connections. **d.** Distribution of distances between functionally connected nodes shows short-, medium- and long-range connections within the ALI-CO. Peak of highest correlated connection distances occurs at approx. 400 μm, medium correlated at approx. 600 μm and lowly correlated at approx. 800 μm. **e.** Schematic diagram of the ALI-CO-mouse spinal cord co-culture with a representative image (right panel) taken 8 days after sectioning. Dorsal (D) and ventral (V) labels denote anatomical coordinates of the explant. **f.** Example fusion of human ALI-CO derived tracts into mouse spinal cord 33 days after initiation of co-culture. Brightfield image (left panel) shows overall positioning of the tissues, with evident axon tracts (arrows) projecting from the ALI-CO (lower right) to the mouse spinal cord (upper left). Staining (right panel) with the human specific marker STEM121 reveals these are ALI-CO derived, and MAP2 staining reveals mouse neurons within the spinal cord and also stains neurons of the human tissue, which is double positive. **g.** Immunofluorescence using a human specific antibody against the presynaptic marker synaptophysin, STEM121, MAP2 and the post-synaptic marker PSD95 and high magnification imaging in the mouse spinal cord region of a mouse spinal cord-ALI-CO co-culture after 32 days reveals mature human-mouse synapses with human SYP juxtaposed to PSD95 along the surface of mouse MAP2+/STEM121-neurons. **h.** Brightfield images of 31-day human-mouse co-culture used for panels i-l. Images show pre- (outlined in blue) and post-(outlined in red) axotomy with a stimulation electrode (yellow dashed outline) placed on axon tracts leading from the organoid. Yellow box: ROI used for muscle contraction traces, white dashed box: region shown in Supplemental Fig. 4e, red dashed line: site of axotomy. **i.** Spontaneous muscle contractions recorded as displacement within the ROI of time lapse imaging of the co-culture shown in h. Recording was made before any manipulation or stimulation of the tissue. **j.** Muscle displacement (contraction) amplitude in response to increasing amplitude of single current pulses (black arrows, 2 mA to 20 mA, 120 μs-long) at 30, 40, 50 and 60 s of the recording (Supplementary Video 11). **k.** Evoked muscle contractions (Supplementary Video 12) pre- (blue) and post- (red) axotomy in response to single current pulses (black arrows, 15 mA, 120 μs-long) applied at 30, 60, 75, 90, 105, 120, 135s during the recording (150 s total duration). **l.** Evoked muscle contractions (Supplementary Video 13) pre- (blue) and post-axotomy (red) in response to 30 s of 1 Hz TTL stimulation with 15 mA current pulses (black hash marks, 120 μs-long). Both spontaneous and evoked contractions were abolished by axotomy (Supplementary Video 14) in k and l, while only a few very low amplitude contractions are seen. **m.** Overlay of evoked muscle contraction waveforms elicited by repeated current pulses (25 mA, 180 μs-long, 0.6 Hz), and waveform average (black), showing the first peak at ~300ms after stimulation. **n.** Distribution of latencies from stimulation to onset of muscle contraction for contractions shown in m. **o.** Evoked muscle contractions to 30 s of repeated current pulses (blue trace, 0.4 Hz, 25 mA, 180 μs-long) in a 21-day human-mouse co-culture (Supplemental Fig. 4i) were abolished by application of 2 uM TTX (orange trace) and restored (green trace) following wash-out with warm media (8x) and 30-minute incubation at 37 °C.

We next examined the functionality of the escaping, extracortical projecting tracts. ALI-COs were co-cultured with sections of the spinal column dissected from embryonic mice in which the associated peripheral nerves and paraspinal muscles were still intact^15^ (Figure 4e). After 2-3 weeks in co-culture, dense axon tracts from the ALI-COs fused with the mouse spinal cord (Figure 4f) and synapses were visible between human projecting axons and mouse spinal motor neurons (Figure 4g). Live imaging of the mouse muscle tissue revealed sporadic concerted muscle contractions with an irregular periodicity (Figure 4h, i, Supplemental Fig. 4c, Supplementary Video 9). Importantly, spontaneous muscle twitches could be observed even in the absence of innervation by human tracts (Supplemental Fig. 4d); however, this activity was less concerted and lower in amplitude than innervated tissue (Supplemental Fig. 4c). In addition, the coordinated, high amplitude contractions of ALI-CO-innervated mouse tissue could be abolished by lesion of the ALI-CO axonal tracts (Supplemental Fig. 4c, Supplementary Video 10) pointing to specificity and dependence on ALI-CO tract innervation.

To test whether ALI-CO extracortical axon tracts could control the mouse paraspinal muscle contractions, we performed extracellular stimulation of the ALI-CO axon tracts and observed muscle response. A single brief stimulation of sufficient intensity of the axon tract was sufficient to elicit a robust muscle contraction (Figure 4j, k, Supplemental Fig. 4e). Evoked muscle contractions were intensity-dependent such that larger stimulation currents increased the amplitude of the muscle contraction (Figure 4j, Supplemental Fig. 4e, Supplementary Video 11), and we could reliably drive the muscle contractions with repeated stimulation (Figure 4k, Supplementary Video 12), and at frequencies up to 1 Hz (Figure 4l, Supplemental Fig. 4e, Supplementary Video 13). The response latency from stimulation to beginning of muscle contraction was approximately 280 ms (Figure 4m, n), which is consistent with a polysynaptic circuit, and suggests electrical stimulation was not activating the muscle directly but relied on the organoid neural tract. Indeed, staining of the responding tissues revealed bundles projecting into the mouse spinal cord (Supplemental Fig. f, g), and lesion of this tract eliminated the ability to evoke muscle contractions (Figure 4h, k,l, Supplemental Fig. 4h, Supplementary Video 14). Finally, application of tetrodotoxin (TTX) to an intact ALI-CO-mouse spinal co-culture prevented evoked muscle contractions, which were restored after TTX washout (Figure 4o, Supplemental Fig. 4i). These findings provide evidence that ALI-CO tracts can produce functional output to an external target.

In conclusion, we find that by culturing cerebral organoids at the ALI, the tissue remains healthy over a very long period of time (we have tested up to 5 months). Importantly, we performed sectioning of organoids after the establishment of the cortical plate as described previously^4^, thus first allowing proper tissue morphogenesis before ALI culture. In this way, the method is highly similar to slice culture typically performed on mid-neurogenesis stage fetal cortex^16^. The axon dynamics observed in ALI-COs is quite striking, and patterns that ensue, including turning behaviors, point to the possibility that there are endogeneous attractive or repulsive cues, similar to *in vivo*. The bundling behaviour further suggests proper tract formation, but importantly, it is distinct from the fasciculation sometimes seen *in vitro* where bundles form indiscriminately between all neighbouring clusters of neurons^17^. Finally, studies with MEAs and explant co-culture point to functional connectivity within and between ALI-COs and external targets. We further show that tractotomy demonstrates specificity and can be used to study the effects of lesion. These experiments are the first to our knowledge to show a functional output from a neural organoid.

## Methods

### Plasmid constructs

The integrating farnesylated GFP construct (pT2-CAG-fGFP, Addgene #108714) and the sleeping beauty transposase plasmid pCAGEN-SB100X as modified from pCMV-SB100^18^ (Addgene #34879) were used as previously described^4^. The integrating farnesylated FusionRed construct was constructed by replacing EGFP in pT2-CAG-fGFP with FusionRed. pT2-CAG-fGFP was linearized using MluI and EcoRI restriction enzymes. The FusioneRed cassette was amplified by PCR from pCi-C-FusionRed-DEST (a gift from Harvey McMahon) using primers: FW 5’-GTGCTGTCTCATCATTTTGGCAAAGAATTCATGGTGAGCGAGCTGATTAAGGAG-3’ and RV 5’-GAGGGTTCAGCTTACTCACGCGTGATTTACCTCCATCACCAGCGC-3’

The cassette was inserted into the pT2-CAG-farnesyl backbone linearized by Gibson assembly.

### Human pluripotent stem cell culture and genome editing

H9 and H1 human embryonic stem cells (hESCs) (obtained from WiCell, and approved for use in this project by the U.K. Stem Cell Bank Steering Committee) were maintained in StemFlex (ThermoFisher, cat. #A3349401) on Matrigel (Corning, cat. #354230) coated plates and passaged twice a week using EDTA. For generation of the fFusionRed line the pCAGEN-SB100X (0.125 μg/ml) plasmid and the transposon donor plasmid pT2-CAG-fFusionRed (0.375 μg/ml) were transfected into H9 hESCs with Lipofectamine Stem (ThermoFisher, cat. #STEM00001). After ~10 days from transfection positive cells were harvested as a pool by fluorescence activated cell sorting on a MoFlo XDP cell sorter (Beckman Coulter).

### Generation of cerebral organoids

Cerebral organoids were generated as previously described according to the enCOR method in order to reliably generate forebrain tissue and a proper cortical plate^4^. Briefly, 18,000 cells were plated with PLGA microfilaments prepared from Vicryl sutures. The original set of media described previously^4^ or alternatively the STEMdiff Cerebral Organoid Kit (Stem Cell Technologies) were used for organoid culture. From day 35 onward the medium was supplemented with 1% dissolved Matrigel basement membrane (Corning, cat. #354234) to achieve establishment of the cortical plate. Between day 40-60 of the protocol plasmids were delivered to the organoids by injection and electroporation in the ventricles. Approximately 1-2 weeks after electroporation the organoids were processed for ALI-CO culture.

### Electroporation of cerebral organoids

Glass microcapillaries (Drummond Scientific, 1-000-0500) were pulled using a P2000 micropipette puller (Suttern Instrument) with the following settings: heat – 550, filament – 5, velocity – 25, delay – 150, pull – 150. The microcapillaries were opened using dissecting scissors to obtain a tip taper of ~8-9 mm. A total of 5 μl of a 320μg/μl plasmid solution (280 ng/μl pT2-CAG-fGFP and 80 ng/μl pCAGEN-SB100) was used for injection and electroporation. Electroporation settings were as previously described^1^.

### Air-Liquid Interface Cerebral Organoid (ALI-CO) culture and live imaging

ALI-CO cultures were prepared using a modified slice culture protocol^19^. Mature organoids (approx. 55-60 days old, in some cases up to 90 days) were harvested using a cut plastic P1000 pipette tip, washed in HBSS without Ca^2+^ and Mg^2+^ (ThermoFisher, cat. #14175095) and embedded in 3% low gelling temperature (LGT) agarose (Sigma, cat. #A9414) at ~40 °C in peel-a-way embedding molds (Sigma, cat. #E6032), typically 1-3 organoids per mold. The agarose blocks were incubated on ice for 10-15 min and processed on a Leica VT1000S vibrating microtome in cold HBSS. 300 μm-thick sections were collected onto Millicell-CM cell culture inserts (Millipore, cat. #PICM0RG50) in 6-well plates and left to equilibrate for one-hour at 37 °C in serum-supplemented slice culture medium (SSSCM): DMEM, 10% FBS, 0.5% (w/v) glucose, supplemented with penicillin-streptomycin and Fungizone. SSSCM was then replaced with serum-free slice culture medium (SFSCM): Neurobasal (Invitrogen, cat. #21103049), 1:50 (v/v) B27+A (Invitrogen, cat. #17504044), 0.5% (w/v) glucose, 1X (v/v) Glutamax supplemented with Antibiotic-Antimycotic (ThermoFisher, cat. #15240062). ALI-CO cultures were maintained in SFSCM at 37 °C and 5% CO_2_ with daily media changes. Media was provided only below the filter insert so that sections stayed at the ALI and were not submerged.

For live imaging, ALI-COs were left to flatten for at least one day before extended imaging on a Zeiss LSM 780 or 710 confocal microscope with incubation chamber set at 7% CO_2_ and 37 °C. Time lapse movies were taken at 10 minute intervals over several hours or days. Image analysis was performed in ImageJ. Temporal projection images were generated from time lapse stacks using the Temporal-Color Code tool in FIJI. Axon growth cone tracing was done using MTrackJ^20^ with manual tracking of growth cones. Data was then exported and plotted as the distance from the start of the track over time. Linear regression was performed on all tracks and the average best fit line calculated for each group (late and early). Orientation analysis was done using the OrientationJ analysis plugin^21^ using Gaussian Gradient setting, with hue determined by the orientation of tracts, brightness determined by coherency, and saturation set at constant. Kymographs were generated using the MultiKymograph tool.

### Histological and immunohistochemical analysis

Organoids and ALI-COs were fixed in 4% PFA for 20 min at room temperature or overnight at 4 °C and washed 3 times in PBS. For cryostat processing, samples were incubated overnight in 30% sucrose. Embedding, cryosectioning and staining were performed as previously described^22^. Whole ALI-COs were stained using a modified protocol – all staining steps were done in permeabilisation buffer (0.25% Triton-X, 4% normal donkey serum in PBS) at 4 °C and their duration was extended as follows: permeabilisation – overnight, primary and secondary antibody incubation – 2 days, wash steps – 8 hours each. Primary antibodies used with corresponding dilutions were: chicken anti-Map2 (Abcam, ab5392, 1:500), mouse anti-Map2 (Chemicon, MAB3418, 1:300), rat anti-Ctip2 (Abcam, ab18465, 1:300), mouse anti-Satb2 (Abcam, ab51502, 1:200), rabbit anti-Cux2 (Abcam ab130395, 1:200), mouse anti-SMI312 (BioLegend, 837904, 1:500), mouse anti-c-Fos (EnCor, MCA-2H2, 1:100), mouse anti-Piccolo (Origene, TA326497, 1:100), mouse anti-STEM121 (Takara, Y40410, 1:500), sheep anti-human-Neuropilin-1 (NRP1) (R&D Systems, AF3870, 1:200), rabbit anti-Homer 1 (Synapti Systems, 160003, 1:100), rabbit anti-PSD95 (Abcam, ab18258, 1:500), mouse anti-human-Synaptophysin (EP10) (ThermoFisher, 14-6525-80, 1:200), goat anti-human-Synaptophysin (R&D Systems, AF5555, 1:100), chicken anti-GFP (ThermoFisher, A10262, 1:500), mouse anti-Bassoon (Enzo, SAP7F407, 1:200), rabbit anti-VGAT (Synaptic Systems, 131013, 1:1000), mouse anti-Calretinin (Swant, 63B, 1:500), rat anti-Somatostatin (Millipore, MAB354, 1:100), mouse anti-GAD67 (Millipore, MAB5406, 1:100).

### Cholera toxin subunit B (CT-B), DiI and emGFP labelling

ALI-COs were visualised on an EVOS FL inverted microscope (ThermoFisher) and using a microinjection capillary <0.2 μl of 1 mg/ml AlexaFluor 647-conjugate CTB (ThermoFisher, C34778) or 1-4 DiI crystals (ThermoFisher, D3911) were delivered to the target region. Similarly, to achieve sparse neuronal labeling, ALI-COs were injected with <0.2 μl of CytoTune emGFP Sendai fluorescence reporter (ThermoFisher, A16519). Two days after CTB and DiI labeling and 5 days after viral emGFP labelling the samples were fixed for histological analyses.

### Whole-cell patch-clamp recordings

Organoid slices were placed in a submerged chamber and continuously superfused at room temperature with artificial cerebrospinal fluid containing (in mM): 119 NaCl, 2.5 KCl, 11 glucose, 26 NaHCO3, 1.25 NaH2PO4, 2.5 CaCl2 and 1.3 MgCl2 (pH 7.4), saturated with 5% CO_2_/95% O_2_. Whole cell patch clamp recordings were performed in current-clamp configuration using an Axon Multiclamp 700B amplifier (Molecular Devices, San Jose, CA) under a Slicescope (Scientifica, Uckfield, UK) equipped with a 40x objective lens (Olympus, Tokyo, Japan) and a WAT-902H analogue camera (Watec, Newburgh, NY). Signals were filtered at 1 kHz and sampled at 10 kHz with a Digidata 1550A (Molecular Devices, San Jose, CA) connected to a computer running pClamp10 (Molecular Devices, San Jose, CA). Pipettes were pulled from borosilicate glass capillaries (1.5 mm OD × 0.86 mm ID; Harvard Apparatus, Holliston, MA) and typical pipette resistance was 12-15 MΩ. Pipettes were filled with artificial intracellular solution containing (in mM): 145 K-gluconate, 5 MgCl2, 0.5 EGTA, 2 Na2ATP, 0.2 Na2GTP, 10 HEPES, adjusted to pH 7.2 with KOH (osmolarity: 280-290 mOsm). Resting membrane potential (RMP) was estimated in current clamp mode immediately after establishing whole cell configuration. Cells with RMP ≤ −50mV were selected for analysis (Table S1). A series of current steps (800 msec) of increasing amplitude (5pA increments) were applied in current clamp mode to determine the frequency-current relationship.

### Organoid cell dissociation for scRNAseq

Organoid tissue slices (two H9 day 53+22, and four H1 day 53+16, three separate ALI-CO preps total) were transferred from the ALI into a conical tube containing Hibernate Medium (Thermo Fisher Scientific, A1247601) plus 1X B-27 Supplement (Thermo Fisher Scientific, 17504044). Gentle dipping helped remove any remaining agarose. The slices were then placed into a 10cm^2^ tissue culture dish and after washing twice with 1X dPBS (Sigma, D8537), transferred into a gentleMACS C Tube (Miltenyi, 130-093-237) containing 2 ml of Accumax (Sigma, A7089) solution. The C Tube was subsequently attached to the gentleMACS Octo Dissociator (Miltenyi) and ran with the recommended cell dissociation settings. The cell suspension was run through a 70μm strainer to remove any residual cell and debris clumps. A small volume of the strained cell suspension was removed for cell counts with the remaining being diluted 4-fold in 1X dPBS and then centrifuged at 200g for 5 minutes. The cell pellet was then resuspended in dPBS containing 0.04% BSA (Sigma, A9418) to give a final concentration of 206 cells/μl. This suspension was kept on ice for 30 minutes until being processed.

### Single-cell RNA-seq library generation and sequencing

Single cell RNA-seq libraries were prepared as instructed by the manufacturer using the 10X Genomics Chromium Single Cell 3’ Library & Gel Bead Kit (10X Genomics, 120237) workflow. The cell suspension (34μl with a total of 7000 cells) was loaded onto a 10X Genomics Chromium Single Cell 3’ Chip with the appropriate amount of Mastermix. The cell capture rate for barcoding varied between 50-75%, resulting in 3500-4400 barcoded cells. The Chromium Controller was run according to the protocol producing the single cell gel beads in emulsion mixture. The reverse transcriptase reaction and subsequent amplification was carried out in a C1000 Touch Thermal Cycler (Biorad), with the cDNA being amplified for 12 cycles. Before sequencing, libraries were quality tested using the 2100 Bioanalyzer Instrument (Agilent) and their concentration was measured by qPCR. Samples were pooled together and sequenced on an Illumina Hiseq 4000 platform.

### Single-cell data analysis

The single-cell RNA sequencing data analysis pipeline was constructed using the CellRanger, Seurat and Monocle software packages. The reads were aligned to the GRCh38 reference genome using STAR in CellRanger 2.1.1. This provided a gene expression matrix of 13,333 cells with a median of 2,320 genes and 29,086 mean reads per cell post-normalization. Quality of reads (exonic and intronic) were analysed by FastQC, showing 87.2% fraction reads in cells. Low quality ends were disregarded during the mapping process. Only reads that were uniquely mapped to the transcriptome were used for UMI counting in CellRanger. The read depth was normalized in the ‘Aggr’ function between the libraries of the samples. UMI (transcript) counts were normalized for each cell to the total counts. The values were multiplied by 10,000 and transformed into log-space. Genes expressed in a minimum of three cells and cells expressing between 200 and 5,000 genes with a maximum of 15% mitochondrial genes were opted in during the filtering process. Further data processing was performed using the Seurat 2.3.0 R package with recommended settings. This yielded a final object of 13,280 cells for the merged libraries, which was then scaled and normalized. Unbiased clustering was achieved by principle component analysis (PCA) using the highly variable genes that were defined by selecting standard deviation as the dispersion function in the ‘FindVariableGenes’ option (bin = 20 for the scaled analysis). PCElbowPLot analysis has guided selecting the maximum number of dimensions assisting the cluster separations process. This was followed by the application of the ‘FindClusters’ function. Clustering was driven by the recommended resolution of 0.16. Clusters were then visualised in 2D and 3D views based on tSNE separation in R. Robustness of cluster identity was determined comparing top differential genes for each cluster with a cut-off at 25% expression frequency within a population, which identified six well-defined clusters. Cell population identities were determined by gene enrichment analysis using cell type, layer, region, dorsoventral position and lobe specific gene sets obtained from databases (Allen Brain Atlas at http://human.brain-map.org) and published work^1,5,11,23–27^. The final cluster identities were then assigned based on the relative proportion of cells expressing the particular reference genes. GO term analysis was performed using the Gene Ontology Consortium online software(http://www.geneontology.org). Gene enrichment was defined by Fisher’s Exact with FDR multiple test correction, and the top 3 biological process annotations for the enriched genes were presented on the basis of the highest fold-enrichment amongst the most significant terms (p < 0.001). To compare the developmental profile of the organoids and the fetal brain, we used the Monocle R-package to derive a pseudotime trajectory of gene expressions from the scRNA-seq data. The raw gene expression matrix of the 12 and 13-week old fetal brain^28^ were processed through the same quality control and filtering process in Seurat as our organoid derived dataset, which served as input to processing in Monocle. Expression levels across the established pseudotime trajectory were observed for the layer markers and visualised in heatmaps comparing data deriving from our organoids and the fetal brain. Gene expression correlation plots were generated in Python Jupyter using matrices generated in Seurat. This analysis was based on the expression of layer marker genes corresponding with distinct developmental cell states.

### Microelectrode array (MEA) recordings

ALI-COs used for electrophysiological recordings were kept in BrainPhys (STEMCELL Technologies, cat. # 05790) supplemented with Neurocult SM1 neuronal supplement (STEMCELL Technologies, cat. # #05793) for a minimum of 12 hours prior to recordings. Extracellular recordings of spontaneous activity in the organoid slices (n=10) were made using a MEA system (MEA1600, Multi Channel Systems). Organoid slices were transferred immediately prior to recording to a 3D grid MEA (60-3DMEA200/12iR-Ti-gr, 60 electrodes, 12 μm diameter, 200 μm spacing, with an internal reference electrode). Media was removed until a thin layer remained allowing the slice to settle on the 3D MEA grid. The temperature was maintained at 37 °C (TC01 controller and TCX-Control software, Multi Channel Systems). During some recordings, tetrodotoxin (1-2 μM in warm media) was applied to the slice using a Pasteur pipette, which was sufficient to block activity (n=3). The signal was sampled at 25 kHz and stored using the 64-channel data acquisition board (MC Card) and MC Rack software (both Multi Channel Systems). Any electrodes with noise fluctuations greater than 50 μV were grounded prior to recording. Data was exported as a binary file to Matlab (MathWorks) for analysis. The raw signal was bandpass filtered (third-order Butterworth, 600-8000 Hz) and a threshold 6 times the standard deviation above the background noise was used to detect extracellular spike waveforms in each channel with a 2 ms refractory period imposed after each detected spike. Correlated spontaneous activity was compared between electrodes using the spike time tiling coefficient^29^ with a synchronicity window (Δt) of 40ms. by translating the publicly available code in C (https://github.com/CCutts/Detecting_pairwise_correlations_in_spike_trains) to Matlab.

### Code availability

All custom code for the MEA analysis in this paper is publicly available at https://github.com/Timothysit/organoids.

### Mouse spinal cord-ALI-CO co-culture

For mouse embryo spinal cord dissection, C57BL/6 pregnant female mice (12.5 days-post-conception) were euthanized and uterine horns harvested by a technician of the MRC LMB animal unit. All steps described from this point on were carried out in ice-cold PBS without Ca^2+^ and Mg^2+^. The uterine horns were cut between implantation sites to separate the embryos. Using precision tweezers (IDEAL-TEK, 5SA) the muscle layer, the decidua and remaining extra-embryonic tissue were removed to isolate individual embryos (E12.5). For isolation of spinal cords together with dorsal root ganglia and paraspinal muscles, 0.15 mm ø dissecting pins (Fine Science Tools, 2600215) were inserted in the head and pelvic-regions to stabilise the embryo. First, the embryo was placed face-down with straddled limbs, the skin overlying the spinal cord was peeled off and the limbs were removed by cutting the embryo along its length in a posterior-to-anterior fasion, at a distance of approximately 1 mm from the midline, on either side of the spinal cord, using a 3 mm cutting edge spring scissors (CohanVannas, 15000-01). The embryo was then repositioned on its side and the internal organs were removed. Last the embryo was placed ventral side up, any remaining undesired tissue was removed including the head and tail.

The dissected mouse spinal cords with surrounding muscle tissue were incubated in ice-cold PBS until embedding. Typically, one organoid and two mouse tissues were first washed in ice-cold HBSS without Ca2+ and Mg2+, then washed and embedded in 3% low-melt agarose at ~40°C. Prior to sectioning and before agarose polymerization, the organoid was positioned centrally at the bottom of the mold on top of a layer of solidified agarose. The mouse tissues were placed flat on either side with their roof plates facing inwards towards the organoid (~1-3 mm away). The organoid was oriented such that its longest axis would be parallel to the spinal cords, thus ensuring the maximum number of slices cut that included both organoid and mouse tissue. The block was oriented on the stage for sectioning along the axial plane of the spinal cords. Sectioning was performed as outlined above. Mouse-organoid co-culture slices were maintained on SFSC medium with daily media changes. After two-three weeks human tracts could be seen innervating mouse spinal cords.

### ALI-CO stimulation and axotomy

Spontaneous contractions of mouse spinal muscles in the organoid-mouse fusions were typically observed after 20-30 days in co-culture using light microscopy and were recorded using a Nikon TE2000 equipped with Andor Neo sCMOS camera using the NIS Elements software for image acquisition. Images were acquired at ~2.3 Hz or ~50 Hz for estimation of latency. Data was acquired as either a .avi or .nd2 file and analyzed using a custom macro in ImageJ (NIH) to calculate the displacement of the tissue upon muscle contraction as a function of time. Muscle contractions evoked by extracellular stimulation of the organoid axonal tracts were elicited using a stainless steel electrode (57100, A-M Systems) connected to a constant current isolated stimulator (0.2-30 mA, 120-180 μs manually- or TTL-triggered pulses, model DS3, Digitimer). When operating the stimulator in TTL-triggered mode, TTL pulses of a given frequency (5 ms duration) were generated using an Arduino running on its internal clock.

For analyses of the latency, we simultaneously recorded the TTL triggers (which trigger the stimulator) and the camera “expose-out” TTL (which indicates when each frame is exposed) using an oscilloscope with a sampling rate of 200kHz (Picoscope 2406B). This allowed us to precisely (±5μs) measure the delay between the first frame of the movie and the previous TTL pulse, from which we could compute the time delay between each frame and the previous stimulation based on the hardware timestamps of each frame and the frequency of the TTL pulses.

Application of tetrodotoxin (TTX, 2 μM) abolished axon tract stimulation evoked muscle contractions after 15 minutes. Washing the fusion with warm media to remove the TTX restored the ability to evoke muscle contractions after 30 minutes in the incubator at 37 °C. For axotomy, the filters were retrieved from the imaging vessels, placed on a plastic support, visualized on an inverted cell culture microscope at 10X magnification and the incision was performed using a microknife (FST, cat. #10316-14). In order to detect the muscle activity from acquired image sequences, the average of the difference between two consecutive frames for the region of interest (ROI) was computed. The resulting temporal signal was then decomposed into the sum of a baseline, stimulation and residual motion. The baseline was estimated as a 1D rolling ball and the stimulation peaks were detected as outliers (12 × SD) from the mean. The overall processing was implemented as a macro for ImageJ that was applied to a selected ROI. The latencies were computed using a Matlab script as the delay between each recorded stimulation TTL pulse and the time when the nearest residual motion went above 1 a.u. as calculated by this motion detector plugin.

### Data Availability Statement

The data that support the findings of this study are included in the supporting material with the paper. Raw data (e.g. raw images and electrophysiological recordings) is available upon request from the corresponding author.

## Acknowledgements

The authors would like to thank members of the Lancaster lab for helpful discussions, and Denis Jabaudon for insightful comments. We also thank members of the LMB mouse facility for help with timed matings and tissue collections, members of the Harvey McMahon laboratory for plasmids, and the LMB light microscopy facility for assistance with imaging. We are grateful to Paul Coupland and Stephane Ballereau (CRUK) for technical assistance and to Marco Galardini and Pedro Beltrao (EBI) for helping with computing resources. Work in the Lancaster laboratory is supported by the Medical Research Council (MC_UP_1201/9). Research in the Lakatos laboratory has been funded by the Medical Research Council (MR/P008658/1), Wellcome Trust (ISSF_<u>RRZC/115_RG89529</u>), Newton Trust (<u>RRZC/115</u>_RG89305), and A.L. is holding an MRC Clinician Scientist Fellowship. Work in the Paulsen lab is supported by grants from the Biotechnology and Biological Sciences Research Council (BBSRC). M.T. receives support from Medical Research Council (MC_UP_1201/2) and the European Research Council (ERC Starting Grant, STG 677029). E.D. receives support from Medical Research Council (file reference number MC_UP_1201/13) and HFSP (CDA).

## Author contributions

S.L.G. planned and performed experiments, analyzed data, and wrote the manuscript. S.B.M. planned and performed MEA and stimulation experiments, analyzed data and wrote the manuscript. G.M.G. performed the organoid cell dissociation and provided quality control for the single-cell preparation, and L.M.D.W. analyzed the RNAseq data. L.M. performed whole-cell electrophysiology experiments and analyzed data under the supervision of M.T. T.S. analyzed MEA electrophysiology data. M.S. generated and maintained organoids and ALI-CO cultures, and analyzed data. J.B. analyzed time-lapse image data. E.D. performed and analyzed time-lapse imaging. O.P. supervised electrophysiology experiments and interpreted data. A.L. planned experiments, analyzed and interpreted data, and supervised G.M.G and L.M.D.W. M.A.L. conceived the project, planned and performed experiments, supervised the project, and wrote the manuscript.

---

**Supplemental Figure 1.**
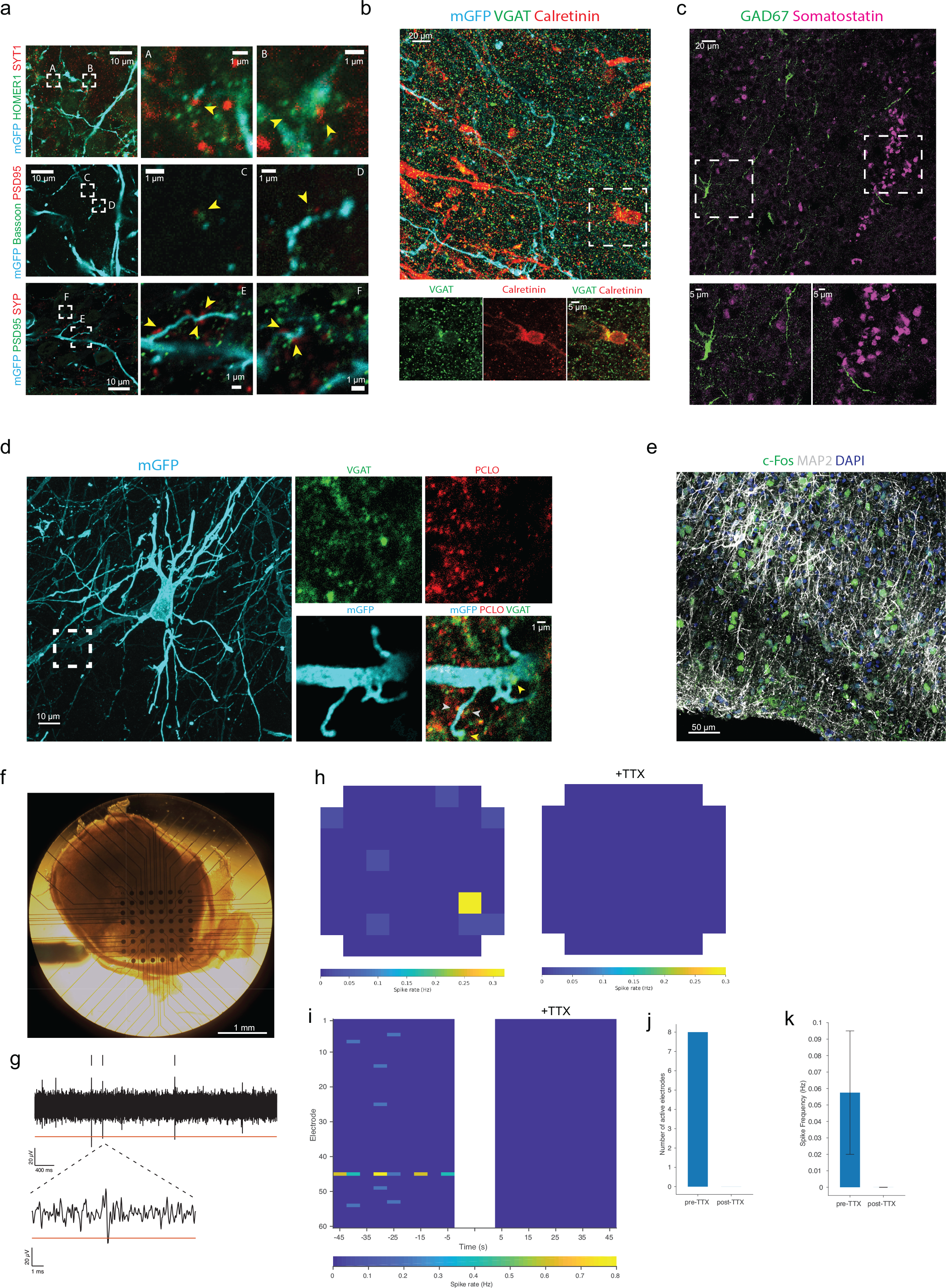
ALI-COs exhibit synapses, interneuron populations, and electrophysiological activity. **a.** ALI-COs from organoids electroporated with fGFP to label neuronal processes harbour mature synapses with juxtaposed pre- (Syt1, Bassoon, Syp) and post- (Homer1, PSD95) synaptic termini. ALI-CO age: 55 + 40 days ALI (95 total) and 89 + 23 (112 total). **b.** Immunostaining for the markers VGAT and calretinin reveal numerous interneurons and VGAT positive puncta suggesting GABA-ergic synapses. ALI-CO age: 57+49 day ALI. **c.** Staining for somatostatin and GAD67 demonstrate the presence of other interneuron types. ALI-CO age: 57+49 day ALI. **d.** ALI-CO electroporated with fGFP reveals a pyramidal neuron with dendritic spines decorated by punctae of the presynaptic structural protein PCLO and the presence of VGAT punctae suggesting GABA-ergic synapses onto the pyramidal neuron. ALI-CO age: 55+40 day ALI. **e.** c-Fos staining for active neurons and MAP2 (dendrites) in a 54+60 day ALI-CO. **f.** Image of an ALI-CO after 44 days at ALI then placed on the 3D multielectrode array (MEA). **g.** Five-second trace from an individual electrode recorded from a 40 + 85 day ALI-CO with threshold (red line) for detected spikes (black hash marks above trace); inset, expanded view of a single spike. **h.** Heatmap of average spike frequency per electrode over 75-second recording before (left) and after (right) application of 2 μM tetrodotoxin (TTX) showing spatial arrangement of spontaneous activity. **i.** Rasterplot of spike frequency (averaged in 5-ms time bins for each electrode) in the ALI-CO 75 seconds before (left) and after (right) TTX illustrating the temporal distribution of spontaneous activity. **j.** Quantification of number of active electrodes before and after TTX application in the ALI-CO recording. **k.** Quantification of average spike frequency (mean+s.e.m.) in active electrodes before and after TTX application.

**Supplemental Figure 2.**
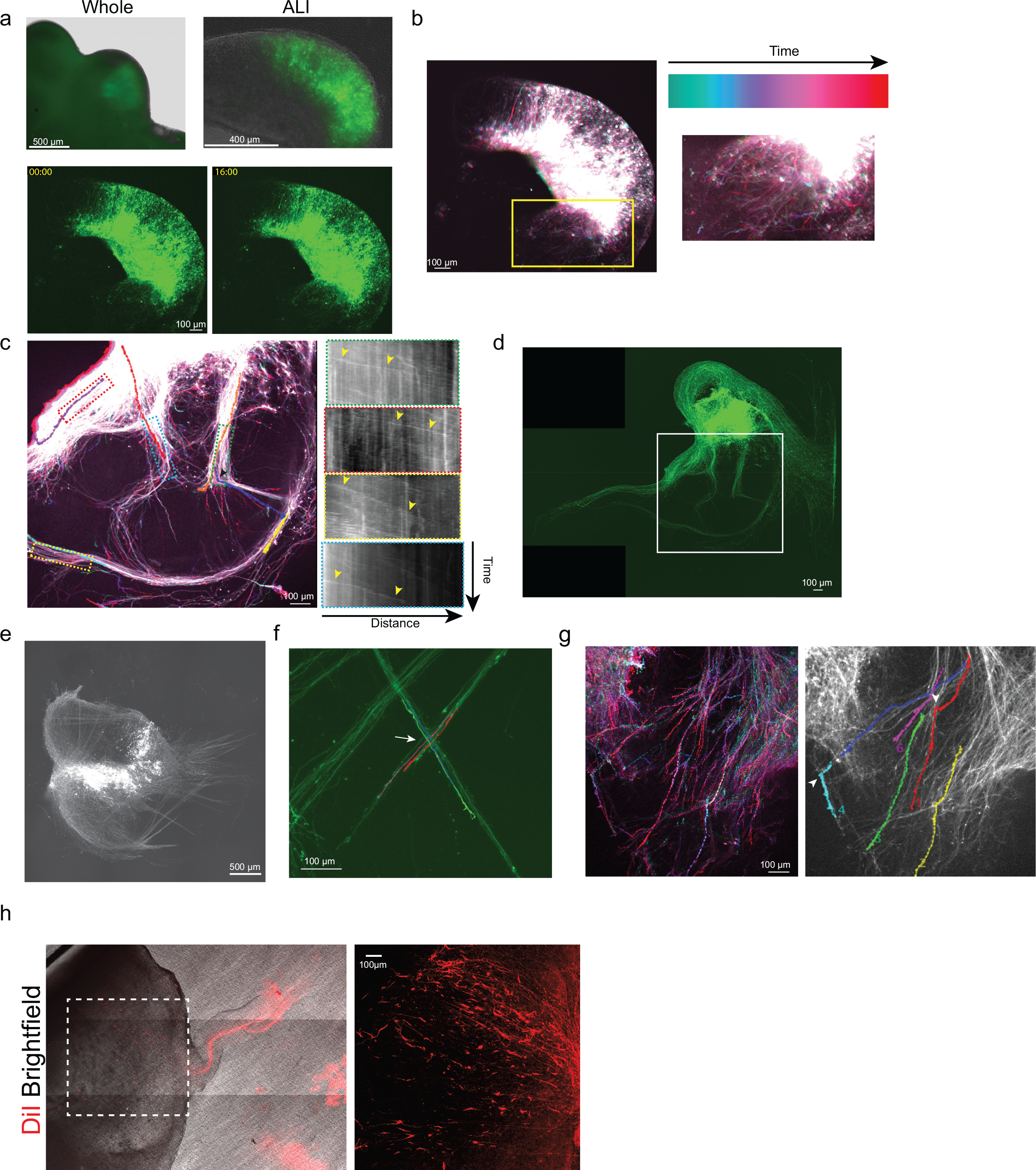
Growth cones within tracts exhibit various behaviors including turning and crossing. **a.** Images of an electroporated organoid before sectioning (upper left), after placement at the ALI (upper right) and stills from longterm live imaging (Supplementary Video 1) of this same tissue (bottom panels). Time stamp is hrs:min. **b.** Temporal projection showing highly disorganized axon outgrowth at this very early stage (2 days ALI). **c.** Temporal projection of a time-lapse (Supplementary Video 5) of a region of the ALI-CO shown in Figure 2e and boxed in d. Overlay includes tracings of individual growth cones (colored lines) and boxed regions used for kymograph analysis (right panels) showing velocities of growth cones (yellow arrowheads). Black arrowhead points to the turning point of an axon. **d.** Image of the same region shown in Figure 2e at an earlier time point of 14 days at the ALI. **e.** Fluorescence image of fGFP+ tracts used for directional image analysis shown in Figure 2f. **f.** Tracing of individual growth cones within intersecting and interdigitated fiber tracts showing crossing behaviour reminiscent of decussation (arrow) (Supplementary Video 6). **g.** Temporal projection (left panel) and growth cone tracing (right panel) showing instances of growth cone turning (arrowheads) (Supplementary Video 7). **h.** Retrograde labelling of an escaping tract by DiI (shown in Figure 2j) and a higher magnification image of the projecting cell bodies (right panel).

**Supplemental Figure 3.**
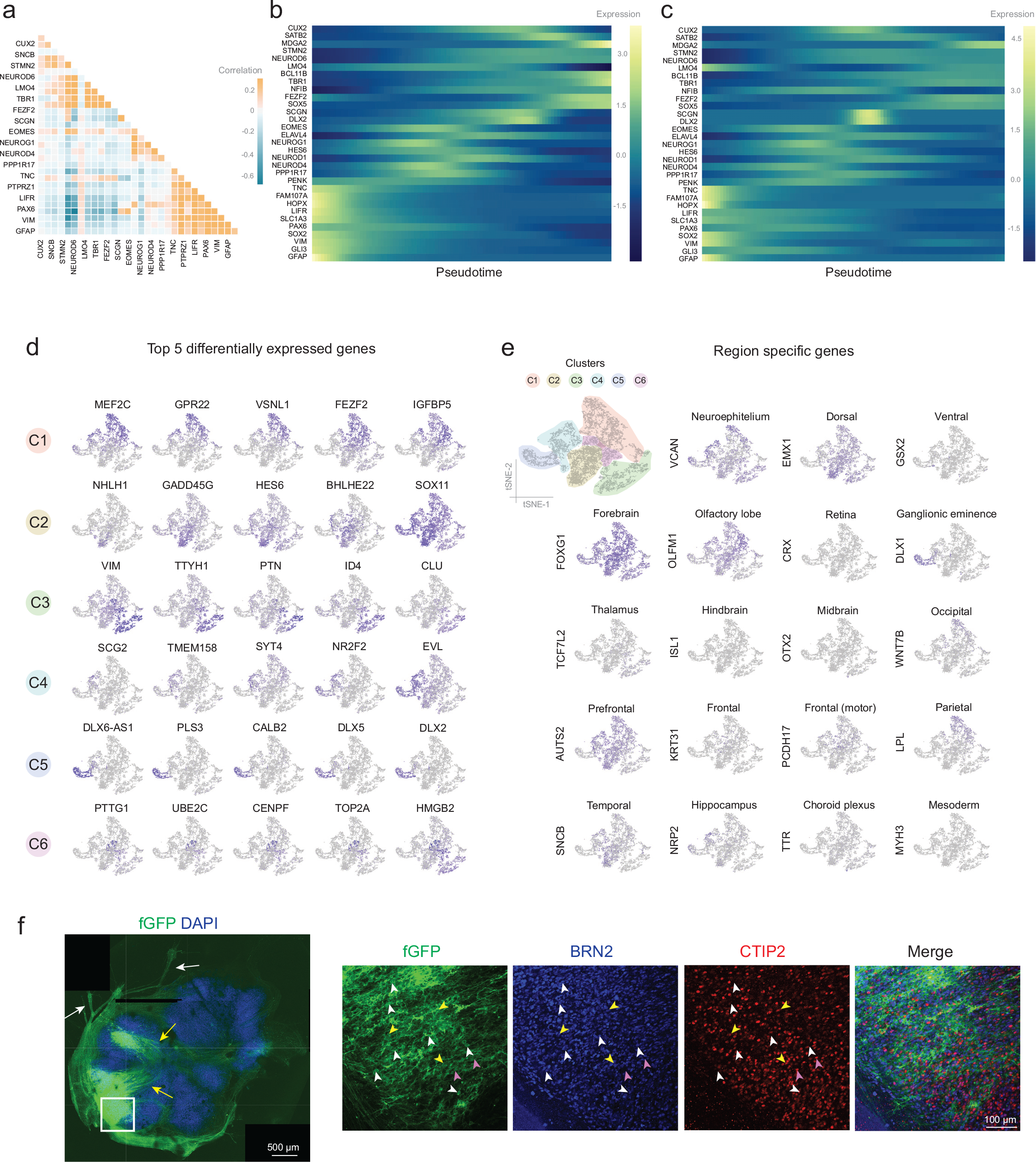
Resemblances of the developmental gene expression profile between the organoids and the fetal brain. **a.** Pearson’s gene expression correlation matrix for developmental layer specific markers in the organoids (blue = negative correlation, yellow = positive correlation). Colour bar represents correlation values. **b.** Heatmaps show normalized layer specific gene expression in the 75-day ALI-COs and **c.** in 12-13 week old human fetal brain cells^28^ ordered by pseudotime. Colour bars reflect logFC gene expression values. **d.** Feature plots for the top 5 differentially expressed genes per cluster in the organoids. **e.** Featuremap shows cell group specific areas in different colours and feature plots demonstrate the distribution of cells expressing region-specific genes across multiple clusters. **f.** Staining for deep layer (CTIP2) and upper layer (BRN2) identities in a fGFP electroporated ALI-CO displaying both internal (yellow arrows) and escaping (white arrows) tract morphologies. Deep layer CTIP2+/BRN2-cells (pink arrowheads) and upper layer CTIP2-/BRN2+ cells (white arrowheads) are visible, as well as some CTIP2+/BRN2+ cells representing a subset of deep layer neurons previously shown to stain for BRN2^30^.

**Supplemental Figure 4.**
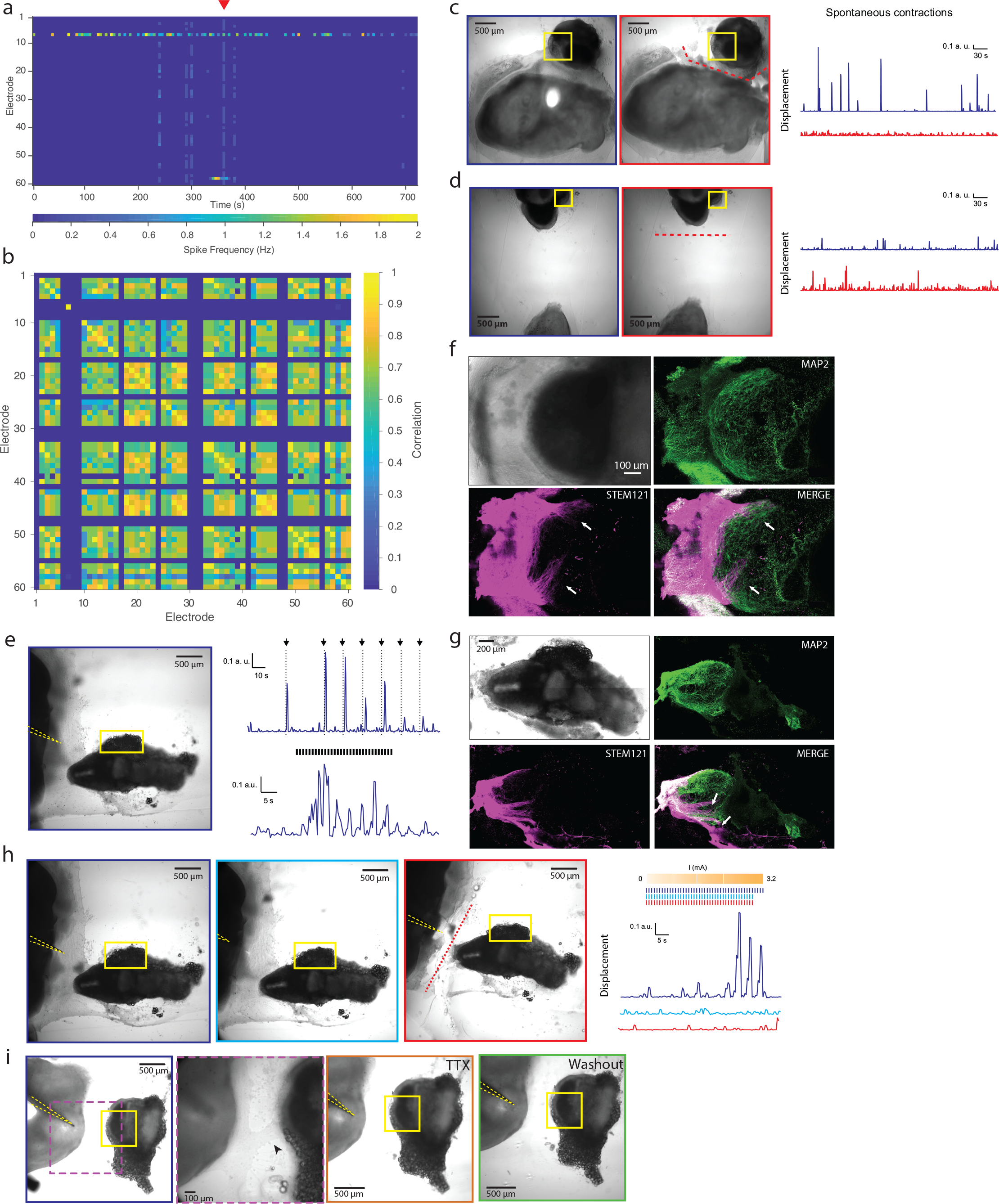
ALI-COs display correlated activity and can elicit muscle contractions in mouse tissue co-cultures. **a.** Rasterplot of spike frequency (averaged in 5-ms time bins for each electrode) during a 6-minute recording. Network burst shown in Figure 4b marked with red triangle. **b.** Correlation matrix shows clusters of synchronous activity. **c.** Human ALI-CO (lower tissue)-mouse spinal (upper right) coculture before (blue box) and after (red box) axotomy (red dash line). Spontaneous contractions (blue trace, Supplementary Video 9) show a decrease in amplitude after axotomy (red trace, Supplementary Video 10). **d.** ALI-CO and mouse co-culture in which fusion has not occurred (blue box) and image after axotomy-style cut was made (red box, red dashed line) near mouse tissue. Spontaneous small amplitude muscle contractions before (blue trace) and after (red trace) cut show similar amplitude and frequency, suggesting that disturbing the mouse tissue through nearby manipulation is not sufficient to eliminate these low-amplitude contractions. Yellow box indicates ROI used for quantification of displacement due to muscle contraction. **e.** Image of 31-day ALI-CO-mouse fusion with a stimulation electrode (yellow dashed line) placed on axon tracts projecting from the organoid. Large amplitude muscle contractions, recorded from ROI (yellow), were evoked by brief current pulses (upper trace, black arrow and dotted line, 3.2 mA, 120 μs-long) applied at 30, 60, 75, 90, 105, 120, 135 s. Repeated stimulation at 1 Hz (lower trace, black hash marks, 3.2 mA, 120 μs-long) could also drive muscle contractions. **f.** Higher magnification image of the mouse spinal section shown in white-dashed box in Figure 4h, stained with MAP2 for mouse neuronal cell bodies and dendrites (green). STEM121 stains human axon bundles (magenta) revealing innervation of the mouse spinal cord (white arrows). **g.** Staining of the same co-culture as in e after axotomy and fixation showing human axon tracts (STEM121+) innervating (arrows) the MAP2+ mouse spinal cord. **h.** Precise electrode placement (yellow dashed line outlining the electrode) determines the ability to evoke muscle contractions in a 31-day mouse spinal cord-ALI-CO co-culture. Left image shows correct electrode placement in the innervating tract with evoked contractions traced at right (blue trace), while electrode displacement from the axon tracts (middle panel, light blue outline), or axotomy (right panel, red outline) abolishes the ability to elicit a response. Red dashed line indicates the site of axotomy. Stimulation before and after electrode displacement and axotomy (hash marks above traces) was done with 1 Hz TTL-stimulated current pulses (120 μs, increasing current amplitude every 10 seconds for 0.2, 0.8, 1.6, 3.2 mA) with the final 1Hz TTL stimulation at 3.2 mAmp lasting 14 s for the control trace (dark blue hash marks). **i.** Confirmation of stimulation electrode placement before (dark blue outline), after TTX application (orange outline), and after wash-out (green outline) for recording shown in Fig. 4o. Yellow boxes denote the ROI used for quantification of muscle contractions. Purple-dashed box with corresponding inset shows the axonal tracts (arrowhead) joining the organoid and the mouse spinal fusion.

Supplemental Video 1.Related to Supplemental Fig.2a and Figure 2d.

Supplemental Video 2.Related to Figure 2b.

Supplemental Video 3.Related to Figure 2c.

Supplemental Video 4.Related to Figure 2d.

Supplemental Video 5.Related to Supplemental Fig.2c.

Supplemental Video 6.Related to Supplemental Fig.2f.

Supplemental Video 7.Related to Supplemental Fig.2g.

Supplemental Video 8.Related to Figure 2g.

Supplemental Video 9.Related to Supplemental Fig.4c pre-axotomy.

Supplemental Video 10. Related to Supplemental Fig. 4c post-axotomy.

Supplemental Video 11. Related to Figure 4j.

Supplemental Video 12. Related to Figure 4k.

Supplemental Video 13. Related to Figure 4l pre-axotomy.

Supplemental Video 14. Related to Figure 4l post-axotomy.

